# An automated workflow to assess completeness and curate GenBank for eDNA metabarcoding: the marine fish assemblage as case study

**DOI:** 10.1101/2022.10.26.513819

**Authors:** Cristina Claver, Oriol Canals, Leire G. de Amézaga, Iñaki Mendibil, Naiara Rodriguez-Ezpeleta

## Abstract

Expectations are high regarding the potential of eDNA metabarcoding for diversity monitoring. To make this approach suitable for this purpose, the completeness and accuracy of reference databases used for taxonomic assignment of eDNA sequences are among the challenges to be tackled. Yet, despite ongoing efforts to increase coverage of reference databases, sequences for key species are lacking, and incorrect records in widely used repositories such as GenBank have been reported. This compromises eDNA metabarcoding studies, especially for high diverse groups such as marine fishes. Here, we have developed a workflow that evaluates the completeness and accuracy of GenBank. For a given combination of species and barcodes a gap analysis is performed, and potentially erroneous sequences are identified. Our gap analysis based on the four most used genes (cytochrome c oxidase subunit 1, 12S rRNA, 16S rRNA and cytochrome b) for fish eDNA metabarcoding found that COI, the universal choice for metazoans, is the gene covering the highest number of Northeast Atlantic marine fishes (70%), while 12S rRNA, the preferred region for fish-targeting studies, only covered about 50% of the species. The presence of too close and too distant barcode sequences as expected by their taxonomic classification confirms presence of erroneous sequences in GenBank that our workflow can detect and eliminate. Comparing taxonomic assignments of real marine eDNA samples with raw and clean reference databases for the most used 12S rRNA barcodes (*teleo* and *MiFish*), we found that both barcodes perform differently, and demonstrated that the application of the database cleaning workflow can result in drastic changes in community composition. Besides providing an automated tool for reference database curation, this study confirms the need to increase 12S rRNA reference sequences for European marine fishes, encourages the use of a multi-marker approach for better community composition assessment, and evidences the dangers of taxonomic assignments by directly querying GenBank.

## Introduction

The reliability of environmental DNA (eDNA) metabarcoding based biomonitoring deeply relies on the accuracy and completeness of reference databases used for taxonomic assignment of eDNA sequences (Virgilio, Backeljau et al. 2010; Richardson, Bengtsson-Palme et al. 2018). eDNA metabarcoding studies can use *ad hoc* built databases, public reference databases or a combination of both. These databases are composed of sequences assigned to species and which cover a specific region of the genome (named barcode) that has enough variability to distinguish among species while being conserved enough to be amplified with universal primers following the principle of the so-called barcoding gap (Hebert, Ratnasingham et al. 2003). Incompleteness of reference databases can lead to false negative detection, leading to for example failure in detecting alien species (Klymus, Marshall et al. 2017); inaccuracy of reference databases can lead to false positive detections, resulting for example in incorrectly reporting species presence (Port, O’Donnell et al. 2016).

Building *ad hoc* reference databases can be performed by barcoding morphologically identified specimens among the ones expected in the area of study, which requires resources and taxonomic expertise and is only possible when the target organismal diversity of the study area is known and low (Cilleros, Valentini et al. 2019; Doble, Hipperson et al. 2020; Milan, Mendes et al. 2020). If the number of expected species in the study area is reduced, sequences of the species of interest can be downloaded and manually curated to remove misannotated sequences, and barcoding of the missing species in public repositories can be performed to complete the reference database (Thomsen, Møller et al. 2016; Collins, Trauzzi et al. 2021; West, Travers et al. 2021). Yet, in some cases, the number of species expected in a region is very large (*e.g.* fishes in a large marine area) and a manual curation of the database is unviable (Leray, Knowlton et al. 2020).

Public reference databases function as open sources of information where researchers submit their sequence data, enhancing reproducibility and transparency (Deiner, Bik et al. 2017; Leray, Knowlton et al. 2020). Some public databases such as BOLD (www.boldsystems.org) have filtering options and analysis tools available for quality controls, such as the trackability of the voucher specimens (Ratnasingham and Hebert 2007), but they only include a few barcodes and often from a reduced number of species. The most complete genetic repository is GenBank (Benson, Cavanaugh et al. 2012), whose free and open submission process is a “double-edged sword” because it leads to unverified record accumulation (Porter and Hajibabaei 2018), some of which result in misannotated sequences (Steinegger and Salzberg 2020). Although the reliability of GenBank for a range of DNA based monitoring applications has been praised (Leray, Knowlton et al. 2019), it has also been contested (Locatelli, McIntyre et al. 2020), and problematic records have been identified in many groups (Steinegger and Salzberg 2020), including fishes (Li, Shen et al. 2018). Effort is being made to identify incorrect records (Leray, Knowlton et al. 2019; Bucklin, Peijnenburg et al. 2021), but their removal takes time because errors are not always reported, much less corrected.

Reference database completeness is especially relevant for diverse groups such as fishes, with about 35,000 described species (Froese and Pauly 2022). Unlike limited and well-described freshwater ecosystems, marine habitats are large and host high ichthyofaunal diversity, which makes manual curation of databases difficult. Marine fishes constitute an important resource globally (FAO 2012), whose management scale monitoring is costly and time consuming with traditional methods. eDNA metabarcoding has arisen as a promising, alternative tool for marine fish monitoring (Gilbey, Carvalho et al. 2021) and this method is being increasingly applied to their study during the last years (Tsuji, Takahara et al. 2019; Fraija-Fernández, Bouquieaux et al. 2020), including invasive species detection (Sepulveda, Nelson et al. 2020), migration pattern discovery (Thalinger, Wolf et al. 2019) or behaviour assessment (Canals, Mendibil et al. 2021). In this context, the availability of curated and complete databases will be foremost for the uptake of eDNA-based approaches in fisheries monitoring.

Fish eDNA metabarcoding studies have been conducted using a variety of barcodes, being the most common ones those based on the mitochondrial cytochrome b (cytb), small (16S) and large (12S) subunit ribosomal RNA (rRNA), and cytochrome c oxidase subunit 1 (COI) genes (Zhang, Zhao et al. 2020). From these, the COI gene is considered standard for animal metabarcoding studies (Hebert, Ratnasingham et al. 2003), which, according to the BOLD public database (Ratnasingham and Hebert 2007) covers a broad range of marine fishes inhabiting European waters (Weigand, Beermann et al. 2019) and for which a fish-specific curated reference database has been developed (Oliveira, Knebelsberger et al. 2016). However, because eDNA extracted from water samples contains traces of many very abundant organisms other than fish, the use of COI metazoan universal primers results in over-amplification of non-target taxa (Collins, Bakker et al. 2019; Fraija-Fernández, Bouquieaux et al. 2020). Thus, fish specific primers have been developed, mostly based on the 12S rRNA gene, such as those amplifying the *teleo* (Valentini, Taberlet et al. 2016) or *MiFish* (Miya, Sato et al. 2015) regions.

Studies using marine water eDNA metabarcoding to assess fish diversity based on 12S rRNA usually perform taxonomic assignment in a variety of ways. Some authors assign taxonomy by directly querying GenBank (e.g. Lamy, Pitz et al. (2021); Sato, Inoue et al. (2021); Zhou, Fan et al. (2022)). Others rely on filtered versions of GenBank containing only the target barcode of the sequences from the species of interest. This filtering can be done either using the reference description (Iwasaki, Fukunaga et al. 2013; Machida, Leray et al. 2017; Arranz, Pearman et al. 2020; Gold, Curd et al. 2021; Mariani, Fernandez et al. 2021; Russo, Maiello et al. 2021; Barco, Kullmann et al. 2022) or based on similarity searches (Heller, Casaletto et al. 2018; Leray, Knowlton et al. 2022). Yet, although the use of these filtered versions is popular in marine fish eDNA metabarcoding studies (Nguyen, Shen et al. 2020; Kawato, Yoshida et al. 2021; Kume, Lavergne et al. 2021; Oka, Doi et al. 2021; Polanco F, Richards et al. 2021), the methods used to extract the sequences are not meant to remove potential contaminations other than those identifiable through their labeling (e.g. Arranz, Pearman et al. (2020)). Finally, other studies attempt to reduce potentially erroneous sequences by visual inspection of phylogenetic trees (e.g. Collins, Bakker et al. (2019); Fraija-Fernández, Bouquieaux et al. (2020); Canals, Mendibil et al. (2021); Collins, Trauzzi et al. (2021)), but this approach is tediuos, not viable for databases composed by a large number of species, and limited by the low phylogenetic resolution of short barcodes (Zhang, Zhao et al. 2020; Polanco F, Richards et al. 2021).

Here, to assist future marine fish eDNA metabarcoding studies, we have developed an automated workflow to i) perform a gap analysis of GenBank for a list of species of interest, ii) create a reference database of specific barcodes for the species of interest, and iii) detect and eliminate the most obvious spurious sequences. As a study case, we have applied this workflow to the fish inhabiting the European Marine Regions. We have assessed the gaps for COI, cytb and 12S rRNA based barcodes, generated a curated reference database for the most widely used (i.e., *teleo* and *MiFish* regions from the 12S rRNA gene) and compared the performance of the taxonomic assignment using the reference database before and after database curation on real, marine eDNA samples. Finally, we contribute to the reference database completeness by barcoding the 12S rRNA sequence of 21 different fish species.

## Materials and methods

A summary of the procedures followed is presented in **Figure S1** and all the scripts used are available at GitHub (https://github.com/rodriguez-ezpeleta/NEA_fish_DB).

### Fish checklist assembly and reference sequence retrieval

The list of fish species present in the northeast Atlantic and adjacent seas was assembled by retrieving the species occurring in the European Marine Regions (Baltic Sea, Barents Sea, Black Sea, Canary Current, Celtic Biscay Shelf, Faroe Plateau, Greenland Sea, Iberian Coastal, Iceland Shelf and Sea, North Sea, Norwegian Sea and Mediterranean Sea), and taxonomy was extracted from World Register of Marine Species (Horton, Kroh et al. 2018). All mitochondrial gene records available in GenBank for the species in the reference list were identified using eUtils (Sayers 2008) and were assigned as belonging to one of the most common genes used in metazoan metabarcoding surveys, i.e., cytochrome oxidase I (COI), cytochrome b (cytb), 12S rRNA (12S) and 16S rRNA (16S), based on their description or, those with ambiguous description, based on BLAST searches (Altschul, Gish et al. 1990) against complete COI, 12S, 16S and/or cytb sequences, respectively; matches were considered if query sequences had ≥ 60% sequence similarity with the complete sequences.

### Barcoding gap analysis

The barcoding gap analysis was carried out for two barcodes of the two most widely used genes in fish metabarcoding studies: *mlCOI* (Leray, Yang et al. 2013) and *folCOI* (Vrijenhoek 1994) from COI, and *teleo* (Valentini, Taberlet et al. 2016) and *MiFish* (Miya, Sato et al. 2015) from 12S rRNA. First, sequences from all COI and 12S rRNA records identified above were downloaded from GenBank. Using mothur (Schloss, Westcott et al. 2009), these sequences were aligned against reference alignments of COI and 12S rRNA reference sequences (previously aligned with MAFFT (Katoh and Standley 2013)), and trimmed to the 12S rRNA and COI regions. Then, complete *folCOI, mlCOI, teleo* and *MiFish* barcode regions were identified using *cutadapt* (Martin 2011), and partial sequences covering at least 90% of the barcode region were identified using mothur with the complete barcode regions as template. Both the complete and partial barcodes were kept for the barcoding gap analysis. Similarity matrices were calculated based on sequence similarity scores obtained by an all-against-all BLAST analysis. Similarity value distributions were visualized in heatmaps for 6 different taxonomic categories: intraspecific (SP), intra-genus (GE), intra-family (FA), intra-order (OR), intra-class (CL) and intra-phylum (PH), and classified into 5 levels so that Level 1 comprises the range of intraspecific similarity values (excluding outliers) and levels 2 to 5 comprises values resulting from dividing uniformly the range of values between the minimum similarity value and the lowest value of Level 1. Pairs with no BLAST-hits among them because not enough coverage or too distant were reported as “No dist”.

### Automated curation of reference databases

To identify potentially erroneous sequences in the database, a series of rules were developed according to how sequences clustered within a given combination of level and taxonomic category focusing on the squares far from the diagonal in heatmaps, which represent sequences that are too similar or too different given their taxonomy. For both inter and intraspecific relationships, a decision tree was created to tag sequences as correct, erroneous or problematic on the basics on how they clustered in a network of “too similar” or “too different” sequences, respectively (**Figure S2**). Using this decision tree, sequences more similar to sequences of other classes or orders than to sequences of the same species, genus or family are tagged as erroneous if there is enough information to conclude which of the sequences is potentially erroneous within the network and tagged as problematic when the information is not enough to resolve it. Networks that are too complicated to resolve by the decision tree can be visually inspected to, combined to other evidence such as blast searches or phylogenetic trees, manually tag the sequences as correct, erroneous or problematic. Erroneous-tagged sequences are automatically removed from the database and erroneous and problematic-tagged sequences are compiled in two independent lists including the reason for their classification as erroneous or problematic. Finally, a curated database is created and outputted in *fasta* and *tax* formats, which are the files required for the posterior use of the database for taxonomic classification.

### Amplicon data generation, bioinformatic processing and analysis

We analysed real marine water samples with the two most used barcodes in fish eDNA metabarcoding studies (i.e., *teleo* and *MiFish*). For that aim, 30 5L-water samples were collected at different locations, time, and depths in the Bay of Biscay (**Figure S3**). Water filtering, DNA extraction and amplification with the *teleo* primer pair (Valentini, Taberlet et al. 2016) were performed as described in Fraija-Fernández, Bouquieaux et al. (2020). For both primer pairs, three replicate PCR amplifications were done per sample in a final volume of 20 μl including 10 μl of KAPA HiFi HotStart ReadyMix (KAPA Biosystems, Wilmington, MA, USA), 0.4 μl of each amplification primer (final concentration of 0.2 μM), 7.2 μl of Milli-Q water, and 2 μl template DNA. The thermocycling profile for PCR amplification with *MiFish* primer pair (Miya, Sato et al. 2015) included 3 min at 95°C; 35 cycles of 20 s at 98°C, 15 s at 60°C and 15 s at 72°C; and finally 5 min at 72°C. Replicate PCR products were combined and purified using AMPure XP beads (Beckman Coulter, California, USA) following manufacturer’s instructions and used as templates for the generation of 12 × 8 dual-indexed amplicons in the second PCR following the “16S Metagenomic Sequence Library Preparation” protocol (Illumina, California, USA) using the Nextera XT Index Kit (Illumina, California, USA). PCR negative controls resulted in no visible amplification in agarose gels. Multiplexed PCR products were purified using the AMPure XP beads, quantified using Quant-iT dsDNA HS assay kit using a Qubit^®^ 2.0 Fluorometer (Life Technologies, California, USA), and adjusted to 4 nM. Then, 5 μl of each sample were pooled, checked for size and concentration using the Agilent 2100 bioanalyzer (Agilent Technologies, California, USA), sequenced using the 2 × 300 paired end protocol on the Illumina MiSeq platform (Illumina, California, USA), and demultiplexed based on their barcode sequences. Quality of demultiplexed reads was verified with FASTQC (Andrews 2010). Primer pairs were removed using *cutadapt* (Martin 2011) allowing a maximum error rate of 20%, and reads longer than 30 nucleotides and containing the two primer sequences were kept and merged using *pear* (Zhang, Kobert et al. 2014) with a minimum overlap of 10 nucleotides for *MiFish* and 20 nucleotides for *teleo*. Pairs with average quality lower than 33 Phred score were removed with *Trimmomatic* (Bolger, Lohse et al. 2014) and those reads shorter than 60 and 140 nucleotides for *teleo* and *Mifish* respectively, not covering the target region or containing ambiguous positions were discarded using *mothur.* Potential chimaeras were detected based on *UCHIME* (Edgar, Haas et al. 2011) and removed. Taxonomy was assigned to unique reads using the Bayesian classifier method (Wang, Garrity et al. 2007) implemented in *mothur* using the *teleo* and *MiFish* databases before and after the automated curation process. Only reads assigned to species level were considered in subsequent steps. Ordination of communities (considering only shared species between both barcodes) was carried out using nonmetric multidimensional scaling (NMDS; *metaMDS* function, *vegan* package version 4.1.1 (Oksanen, Blanchet et al. 2013) analyses based on Bray-Curtis dissimilarities *(vegdist* function, *vegan* package). ANOSIM (analysis of similarity; Clarke (1993)) was used to test if samples were grouped according to the factor barcode (anosim function, vegan package).

### Generation of 12S rRNA sequences

Fin and muscle tissue samples from morphologically identified specimens (**Table S1**) were obtained during the 2020 SUMMER survey in the Western Mediterranean Sea (Balearic Islands, Alborán Sea, Gulf of Cadiz and Atlantic Ocean) and from fishing vessels landing in the port of Ondarroa (Basque Country, Spain). For each sample, genomic DNA was extracted from muscle tissue or fin using the Wizard Genomic DNA Purification kit (Promega, WI, USA) following manufacturer’s instructions for ‘Isolating Genomic DNA from Tissue Culture Cells and Animal Tissue’. Extracted DNA was resuspended in Milli-Q water and its concentration was determined with NANODROP (Thermo Scientific™). The extracted DNA was then amplified using the *MarineFish* primer pair (Jin, Zhao et al. 2013), a 900-1,100 bp-long 12S rRNA region covering both *teleo* and *MiFish* regions, by mixing 10 μL of 2X PCR Master Mix (Fisher Scientific), 0.4 μL of each primer, 2 μL DNA template (1-20 ng), and 7.2 μL of Nuclease Free water, and using the following amplification conditions: 95°C for 3 min; 35 cycles of denaturation at 95°C for 30 s, annealing at 56°C for 30 s, and extension at 72°C for 75 s; and final extension at 72°C for 10 min. The PCR products were migrated in a 2% agarose gel in TAE buffer and purified using ILUSTRA EXOSTAR1-Step (Cytiva) following manufacturer’s conditions and sent for Sanger sequencing. Forward and reverse sequences were merged and SeqTrace software (Stucky 2012) was used for quality filtering (minimum confidence score 30). Sequences were submitted to GenBank (accession numbers available in **Table S1**) and added to the above generated *teleo/MiFish* reference databases.

## Results and discussion

### Database completeness for the most used fish eDNA metabarcoding markers

The list of Northeast Atlantic and Mediterranean marine fishes compiled included 1791 species: 1603 Actinopterygii, 174 Elasmobranchii, 8 Holocephali, 4 Petromyzonti and 2 Myxini (**Table S2**). In total, 1277, 1067, 1047 and 898 fish species have COI, 12S, 16S and cytb gene records available, respectively, including 42,115 COI, 27,546 cytb, 8,542 16S and 6,820 12S sequences (**Figure 1a**). The COI gene is the one with the highest number of sequences and species coverage (70%) and cytb, despite having the second highest number of sequences, is the one with the lowest species coverage (50%). This is due to a high number of cytb records belonging to a small number of species (*e.g*., Atlantic cod *Gadus morhua*, European anchovy *Engraulis encrasicolus*, or milkfish *Chanos chanos*). 12S and 16S rRNA exhibit similar species coverage values (about 60%).

**FIGURE 1.**
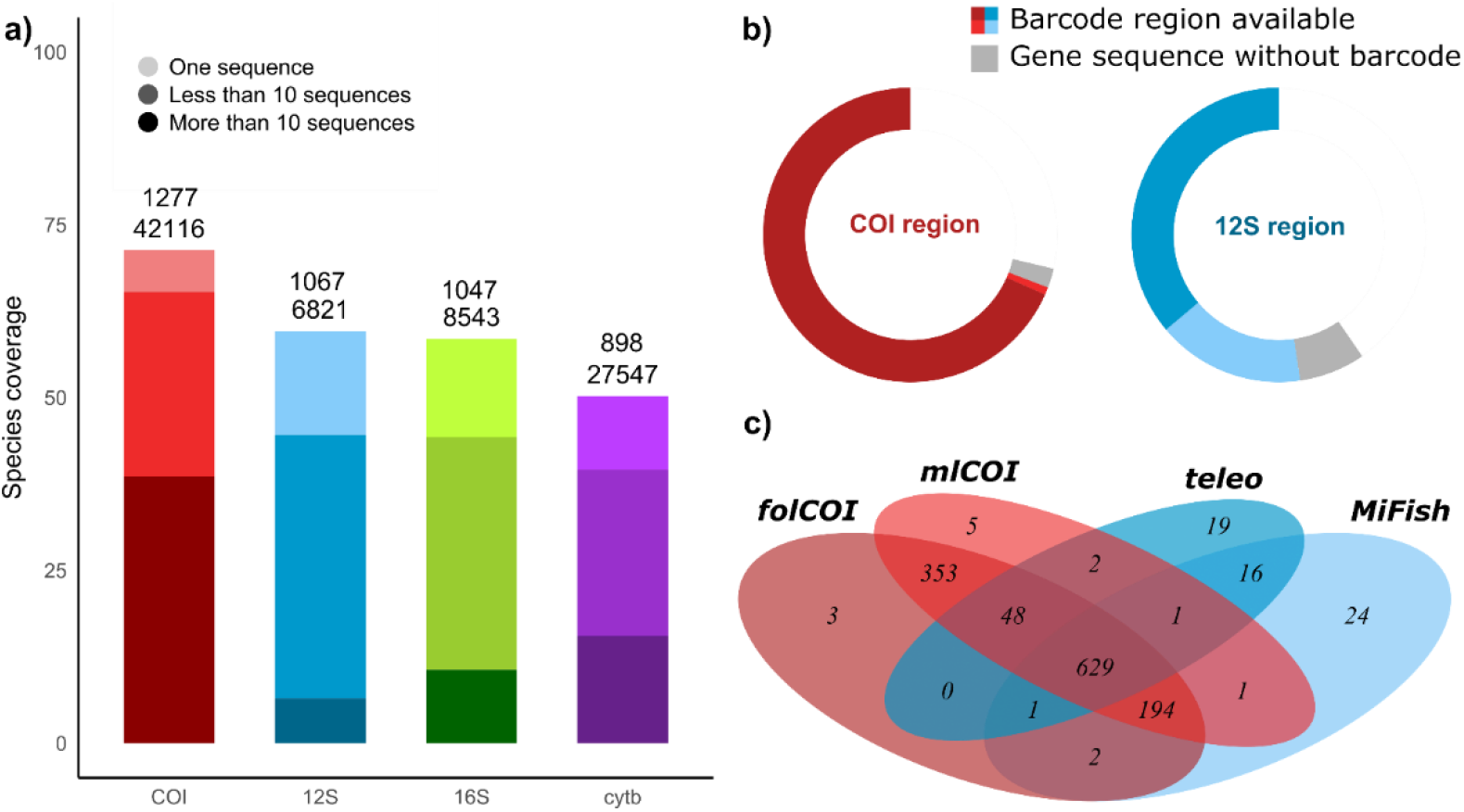
Reference database gap analysis. **a)** Cumulative coverage (%) of European marine fishes for each gene. Numbers on bars indicate the number of species for which there are sequences available (above) and the total number of sequences available in GenBank (below) for each gene. **b)** Barcode availability for COI and 12S gene markers. Dark red and dark blue represent portion of species with both barcodes available (i.e., *mlCOI* and *folCOI* for COI and *Mifish* and *teleo* for 12S). Light red and light blue represent portion of species with only one of the two barcodes available (i.e., *mlCOI* or *folCOI* for COI and *Mifish* or *teleo* for 12S). **c)** Venn Diagram showing the number of species with available references for *mlCOI, folCOI, teleo*, and *MiFish* barcodes.

COI barcodes have the highest, and similar among them, species coverage (>70%), whereas 12S rRNA barcodes cover between 40% (*teleo*) and 48% (*MiFish*) of the species (**Figure 1b**). Recent studies have shown that the COI universal barcodes, despite considered the standard region for barcoding (Hebert, Ratnasingham et al. 2003) and being abundant in public databases (Porter and Hajibabaei 2018), are not effective for fish eDNA metabarcoding as they mostly amplify non-target taxa (Collins, Bakker et al. 2019; Fraija-Fernández, Bouquieaux et al. 2020), and barcodes from other genes such as 12S rRNA have demonstrated to perform better (McClenaghan, Fahner et al. 2020; Zhang, Zhao et al. 2020) even with less species coverage in reference databases (Collins, Bakker et al. 2019). Here, we highlight that the 12S rRNA gene, although being the most used region for fishes, is only sequenced for half of the fish species inhabiting European marine waters. Further, the actual number of species available for specific barcodes (i.e., *teleo* and *MiFish*) is even lower, due to *MiFish* and *teleo* barcodes being non-overlapping—so that the existence of a 12S rRNA sequence for a given species does not imply that both regions are covered, as observed for 45% of the species **(Figure 1c)**. To contribute to complete the 12S rRNA barcode reference database, required for present and future eDNA-based fish monitoring, we have sequenced the *teleo* and *Mifish* regions of 21 species, from which 5 and 16 had none or only one of the barcodes available at the time of submission (**Table S1**).

### Using the barcoding gap principle for potential error detection in reference databases

The so-called barcoding gap relies on the principle that the largest the difference between intraspecific and interspecific genetic distances the more accurate the taxonomic classification (Hebert, Stoeckle et al. 2004). For fishes, the barcoding gap has been examined for large (500- 900bp) mitochondrial regions (Cawthorn, Steinman et al. 2012; Li, Shen et al. 2018), but eDNA metabarcoding usually relies on shorter DNA fragments (≈ 60-170 bp), for which the barcoding gap requires further examination. We assessed the barcoding gap of two COI and two 12S rRNA barcodes and examined the number of pairs of sequences that do not follow the barcoding gap principle, that is, those that are in level 1 (see Methods) but belong to different species, or those that are in higher levels while being taxonomically close. Distance matrices resulted in more than half a billion sequence pair comparisons for both COI barcodes and about 8 and 5 million pair comparisons for *Mifish* and *teleo*, respectively. In all barcodes, the average pairwise similarity decreases as sequences belong to more distant taxonomic categories (**Figure 2a**), but an unexpected number of outliers representing low similarity in pairs of sequences belonging to the same species and high similarity in sequences belonging to taxonomically distant species are noticeable. The categorization of distance ranges in levels (**Table S3**) allows to quantify the number of pairs that do not behave as expected according to the barcoding gap principle (at the most top-right and bottom-left squares of **Figure 2b**). These sequences could be potential errors or contaminations in the reference database, which are known to be present in GenBank (Steinegger and Salzberg 2020), including in fishes (Li, Shen et al. 2018). For these outlier pairs, defining a strategy to identify the erroneous sequence should be feasible considering the high taxonomic distance between them. Cases closer to the diagonal of the heatmap are more complex and make it specially challenging to identify whether sequences are truly erroneous or whether natural reasons make the pair be out of the diagonal. For example, low taxonomic discrimination by the 12S gene has been reported within fish genera (e.g. *Sebastes, Anarhichas*) and families (e.g. *Gadidae, Cyprinidae, Istiophoridae*) (Johnstone, Marshall et al. 2007; Thomsen, Møller et al. 2016; Gold, Curd et al. 2021), which could make sequences appear more similar than expected according to taxonomy. Similarly, biological phenomena such as inter-specific introgression could make sequences from the same species appear more distant than expected and species from different species closer than expected (Viñas and Tudela 2009).

**FIGURE 2.**
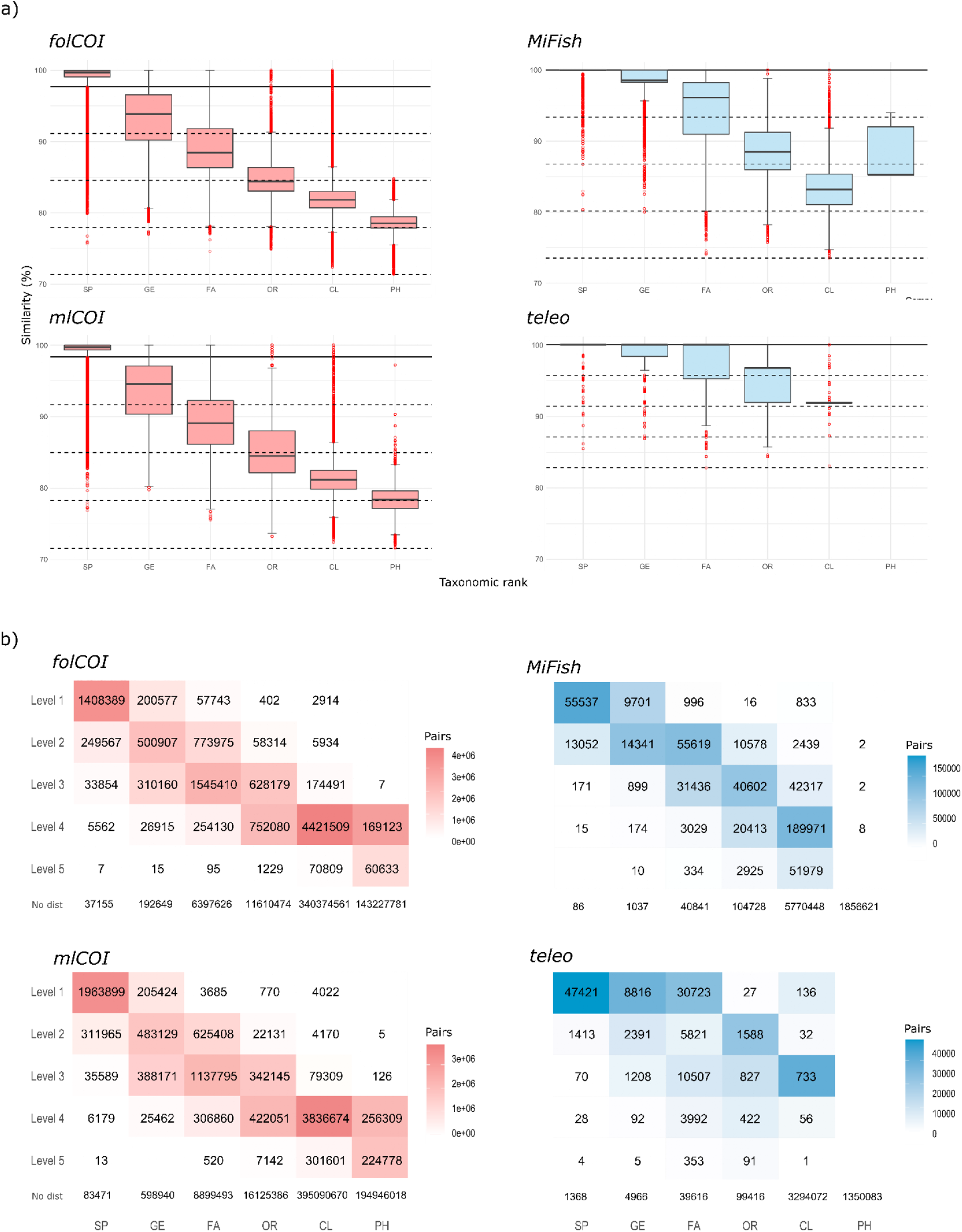
Barcoding gap analysis. a) Boxplots depicting sequence pair distances (%) for intraspecific (SP) and interspecific divergences at different taxonomic levels (GE, FA, OR, CL, PH). Levels are indicated with dashed horizontal lines (note that the values vary for each barcode). Outliers are represented with red dots. b) Heatmaps representing number of pairs within different divergence levels.

### Towards an automated database curation procedure

To date, there is no standard protocol for reference database curation. The most accurate approach for manual curation is the use of phylogenetic trees, which allows detailed inspection for erroneous sequence detection (Collins, Bakker et al. 2019; Leray, Knowlton et al. 2019; Collins, Trauzzi et al. 2021). However, manual inspection of phylogenetic trees is not viable for large databases and has limitations for short and unequal length sequences. Here, we explore an alternative solution and propose an automatized workflow for spurious sequence detection based on network analysis. Briefly, the approach considers spurious sequences those that are more similar to sequences from other species than to sequences of their own (labelled) species. We have focused on 12S rRNA barcodes for being the preferred region for fish eDNA metabarcoding, yet the method can be used for any barcode.

Based on the heatmaps of the previous section (**Figure 2b**), we set the levels to be analyzed. We focused on the extremes, selecting levels 3 to 5 for intraspecific relationships (i.e., too distant pairs) and levels 1 and 2 for intra-phylum, intra-class, and intra-order interspecific relationships (i.e., too similar pairs). For each chosen taxonomic classification and level, independent networks were created, and the decision tree method developed (**Figure S2**) was applied to the resulting clusters of sequences to identify potentially erroneous and problematic sequences. Notably, our approach detected spurious sequences for both 12S rRNA barcodes (summarized in **Tables S4-5)**.

According to the decision tree, when clusters are formed by a central sequence, that central sequence is evaluated. For example, in intraspecific relationships (**Figure S2b**), one central sequence of *Alburnus alburnus* was more similar to *Phoxinus phoxinus* sequences than to other sequences of *A. alburnus* for both *teleo* and *MiFish* (**Figure 3a-b**) and was classified as potentially erroneous. For interspecific relationships concerning central sequences, for example, one sequence of *Carcharodon carcharias* was identical to *Cetorhinus maximus* sequences **(Figure 3-cd**) while showing low similarity to other *C. carcharias* sequences and was also tagged as potentially erroneous. When there is no central sequence, all sequences of the cluster are analysed one by one, such as in **Figure 3e**. In our data, we found *Engraulis encrasicolus* and *Istiophorus albicans* sequences being more similar than expected. Since all sequences of *E. encrasicolus* (including those not represented in the cluster) were very similar to each other and located in Levels 1-2 in the heatmap, (i.e., within expected intraspecific distances), they were thus labelled as correct. For *I. albicans*, there were no other sequences in the database, but *I. albicans* sequences were more similar to *Engraulis* sequences that to other sequences from the *Istiophorus* genus; thus, the *I. albicans* sequences were labelled as erroneous. Interestingly, although present in both the *teleo* and *MiFish* reference databases, *I. albicans* sequences were only labelled as erroneous in the *teleo* database. In the *MiFish* database, *I. albicans* sequences were more similar to other *Istiophorus* sequences than to sequences belonging to the genus *Engraulis*. This can be explained by the formation of chimeras between the target species and other species during the barcoding process, either in the PCR or assembly steps (Edgar 2016). Noteworthy, our method was able to detect this particularly challenging but existing problem in genetic databases.

**FIGURE 3.**
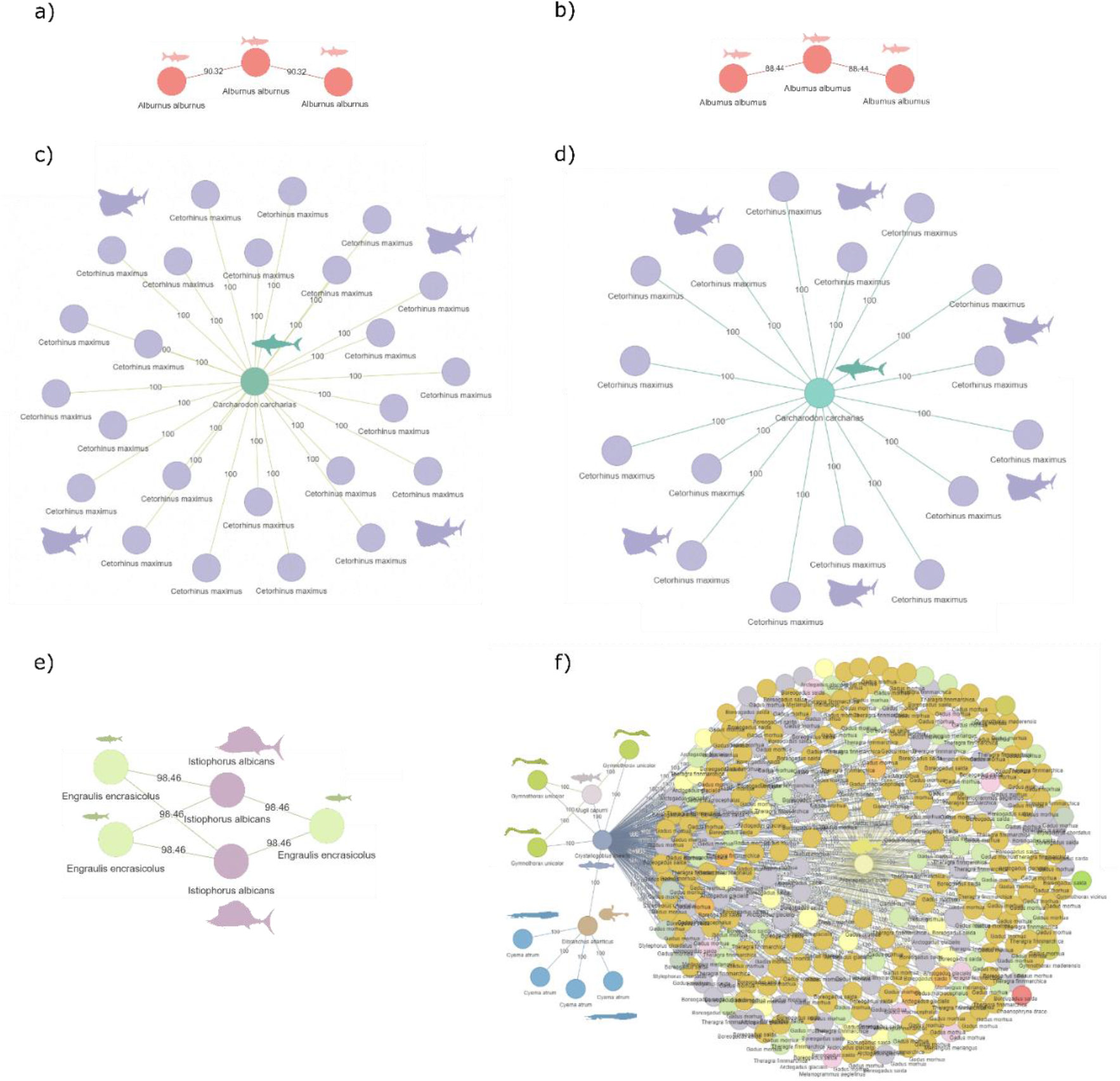
Examples of developed networks for teleo (left) and MiFish (right) databases. a-b) Intraspecific distance analysis of the common bleak Alburnus alburnus sequences of a) MiFish region in level 3 and b) teleo region in level 4. c-d) Interspecific (intra-order level) distance analysis of one sequence of the great white shark Carcharodon carcharias and sequences of the barking shark Cetorhinus maximus in level 1 of c) MiFish and d) teleo sequences. e) Complex structure network formed by two species. f) Complex multi-species structure network of interspecific (intra-class level) comparisons in level 1.

Complex structures formed by more than two species without a central sequence were not approached by our automated analysis, and they should instead be inspected manually. For instance, in **Figure 3f** we can rapidly identify two non-gadoid sequences (belonging to *Argyropelecus gigas* and *Crystallogobius linearis*) that are identical to many Gadidae sequences. In this case, since *Crystallogobius linearis* sequence is more similar to sequences of the Gadidae family than to other sequences of the same species, it would be classified as erroneous. On the other hand, the sequence of *Argyropelecus gigas* would be classified as problematic due to lack of information to compare with in the database because there are no more sequences for *Argyropelecus gigas* and no intra-genus relationships are available.

Starting from any given reference database, our workflow performs a quick screening to detect erroneously labelled sequences and flag problematic ones; additionally, it provides the networks for the sequences that did not result in a clear diagnostic due to the complexity of the distance relationships so that they can be manually inspected. Thus, this workflow is a significant step in automatically improving GenBank based reference databases for diverse taxa. We acknowledge that the tool assumes a linear relationship between similarity and taxonomic relatedness, which is not always fulfilled by real sequences. Yet, this assumption ensures the detection of errors in the extreme cases, and, moreover, the tool allows to modify taxonomic ranges and distance levels to be included in the analyses so that less extreme cases can also be inspected. Thus, this workflow not only allows to retrieve the barcode sequences corresponding to a given list of species, but performs a first screening of spurious sequences, allowing to eliminate, flag or further inspect them.

### Performance of raw and curated reference databases

For the 30 samples included in this study, we obtained 2,274,886 and 1,462,841 *Mifish* and *teleo* reads, respectively (**Table S6-7**), from which ≈90% were assigned to the species level. For *teleo*, the Atlantic sailfish *Istiophorus albicans* represented 35% of the reads when using the raw database. Because *I. albicans* sequences were labelled as erroneous by our automated workflow, reads previously assigned to sailfish using the raw database were classified as anchovy with the clean database, leading to more coherent results across barcodes (**Figure 4**). The fact that no reads were classified as *I. albicans* with the *MiFish* raw database supports the chimeric structure of the sequence, being some regions truly from *I. albicans* and others from *Engraulis*. Contaminant amplification and data entry error cases in GenBank have been reported previously (Leray, Knowlton et al. 2019). Also, misidentifications of sampled specimens can occur, especially when referring to species with morphological similarities (Lyon, Tonkin et al. 2018), rare species, or individuals derived from fishery vessels (Kirsch, Day et al. 2018; Figueiredo, Maia et al. 2020). This can be due to lack of taxonomists (Buyck 1999) or to rapid classification onboard based on the most likely species (FAO 2004). Even if the voucher specimen is correctly identified, additional issues can occur downstream the sample processing and analysis. For instance, contaminant DNA of *Homo sapiens* (Kryukov and Imanishi 2016), bacteria (Strong, Xu et al. 2014), or species in previously extracted samples could result in erroneously labelled sequences in the database.

**FIGURE 4.**
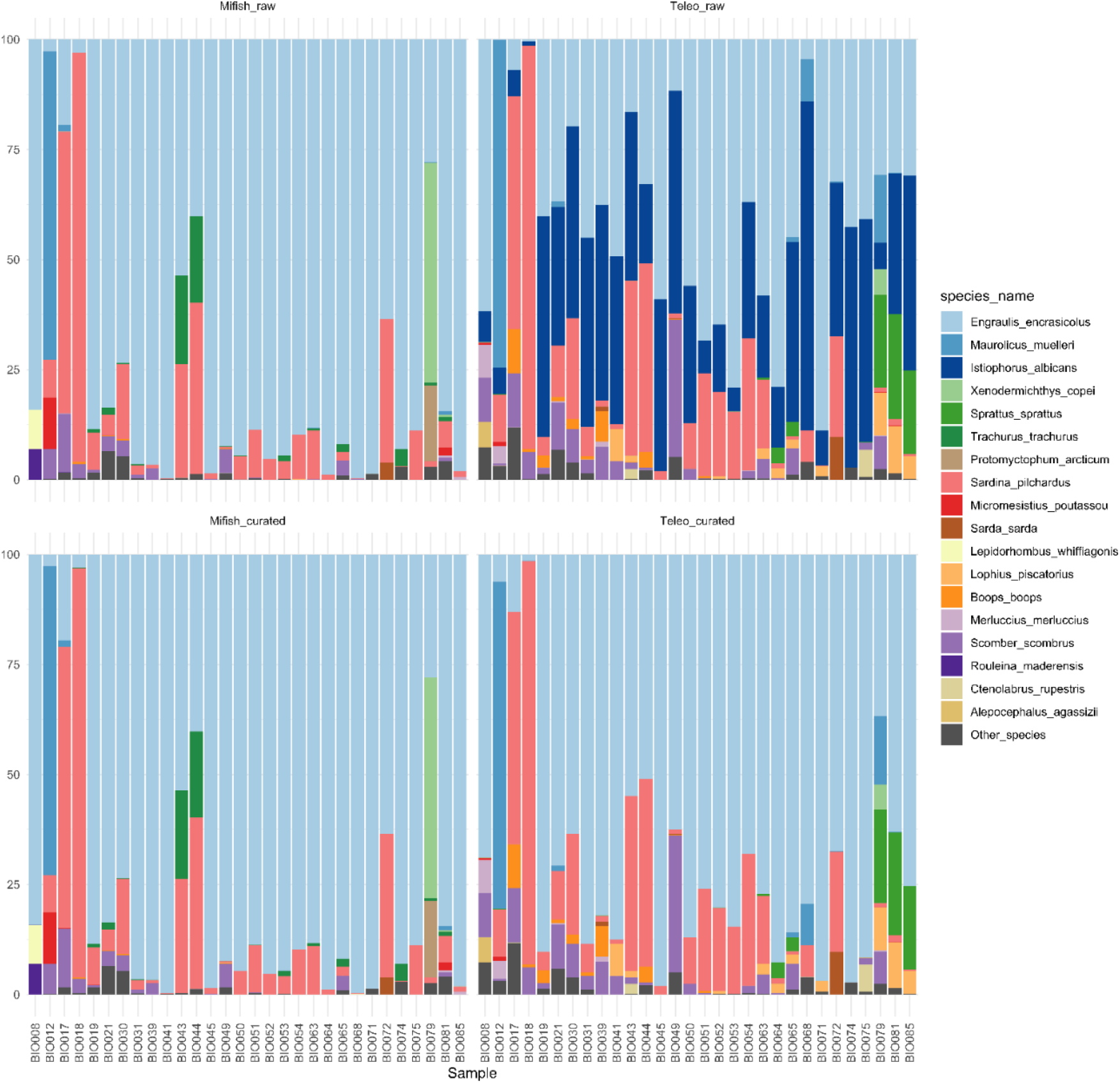
Barplots showing the read relative abundance for the 17 most abundant species using MiFish (left) and teleo (right) barcodes, performing the taxonomic assignment against raw (top) and curated (bottom) reference databases. Less abundant species are merged into “Other_species”.

It is noteworthy to remark that database curation substantially changed the taxonomic assignment of *teleo* reads, which highlights the importance of caution and critical reasoning when analysing metabarcoding data to avoid wrong interpretations or misunderstandings. Although minor differences were observed between the taxonomic assignments of *MiFish* reads using raw and curated versions of the database, potential erroneous sequences belonging to species not detected in the study were also identified, which may be problematic in studies involving other fish assemblages.

**FIGURE 5.**
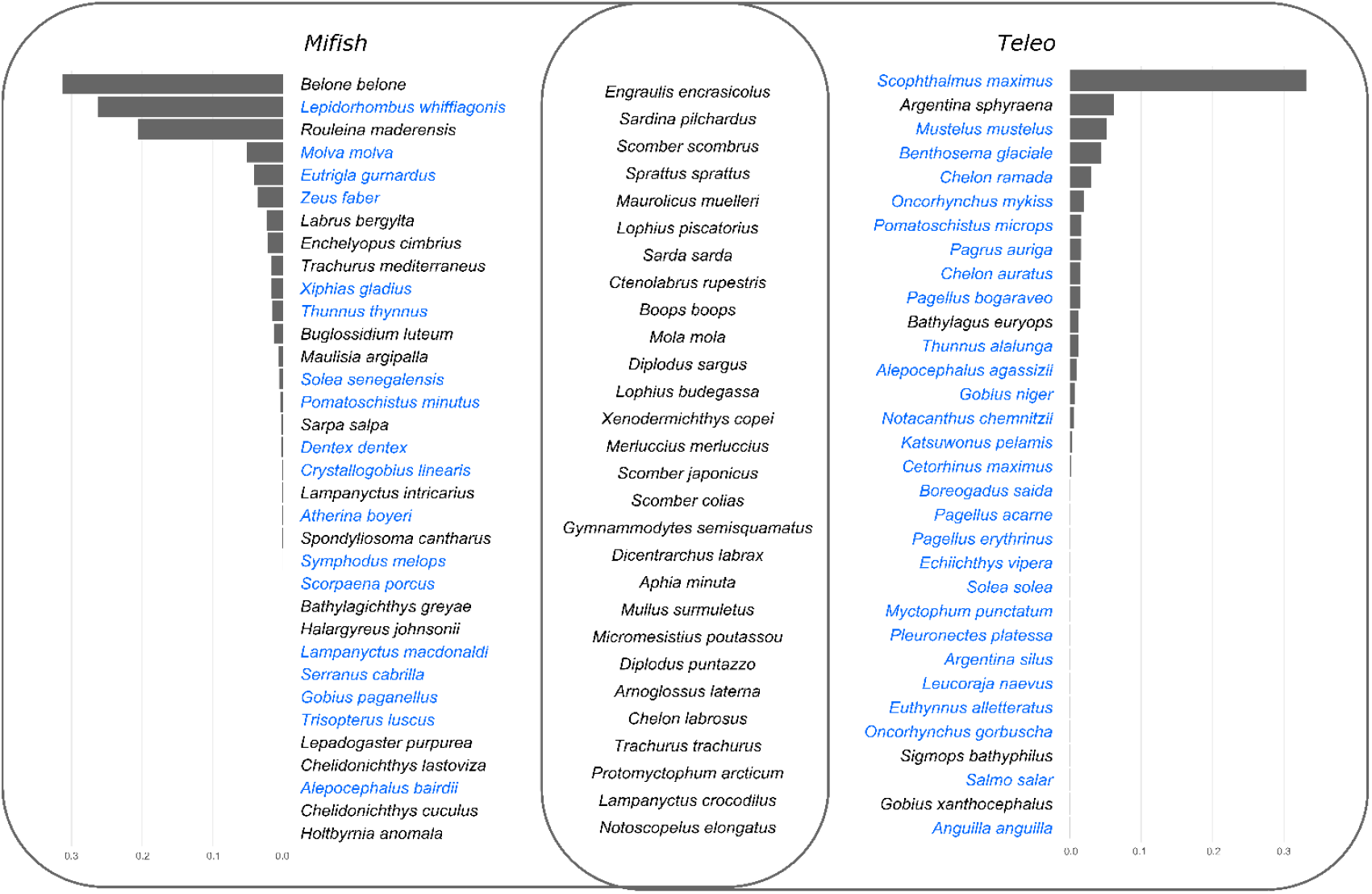
Venn diagram representing the species identified in this study. Barplots indicate the relative abundance (%) of species detected with only one barcode. Species highlighted in blue also have representative sequence in the reference database of the other barcode, but were not detected.

We would also like to point out the differences in fish diversity retrieved by each barcode. A total of 94 species were identified in the study, from which only 30% were detected by both barcodes. The number of species only detected by one of the barcodes was quite high (66 species) (**Figure 5**), suggesting that a multi-marker approach, i.e., the combination of at least two barcodes, is desirable to cover as much diversity as possible, as previously reported (Polanco F, Richards et al. 2021; Kumar, Reaume et al. 2022). Finally, it is noteworthy to mention that we observed a positive correlation between the relative read abundance of most of the species detected by both barcodes (**Figure 6a**), and that the barcode was not supported to be the main factor determining the fish community composition of the samples (**Figure 6b**) (ANOSIM test, R: 0.047, p-value: 0.0161).

**Figure 6.**
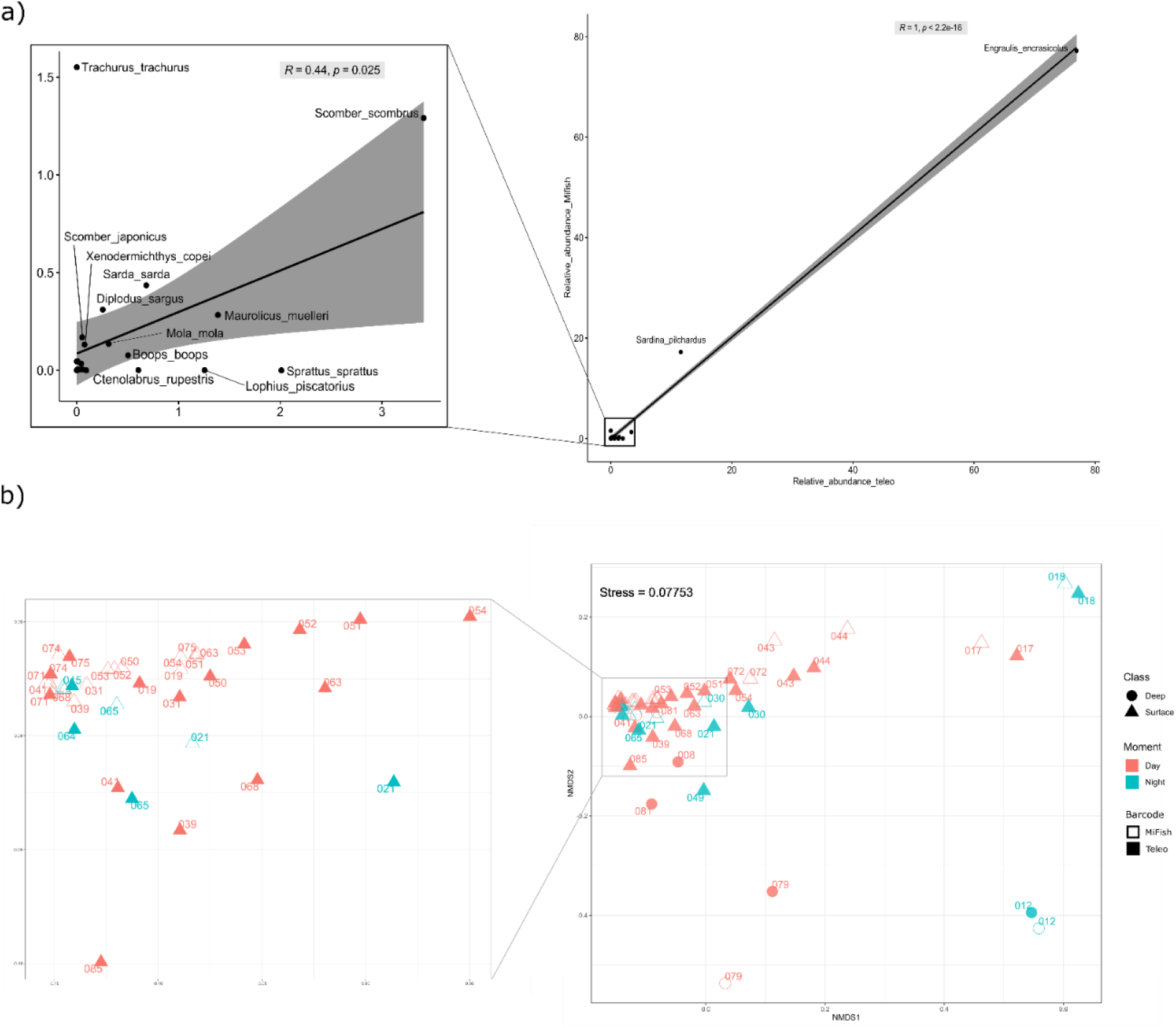
a) Relationship between relative abundance of reads of shared species between Mifish and teleo barcodes. Shaded area represents the 95% confidence interval of the linear regression. b) Non-metric multidimensional scaling (NMDS) for shared species using the two markers.

## Conclusions

Starting from a list of species of interest, our automated workflow performs a gap analysis and downloads and curates a reference database for the barcodes of interest. Applying the method to the marine fish case study, we have detected erroneous sequences in *teleo* and *MiFish* reference databases, which, if not corrected, result in erroneous taxonomic assignments in real marine eDNA samples. This newly developed workflow, which can be applied to any mitochondrial barcode and set of species, and results derived from it constitute a step ahead for increasing the completeness and accuracy of reference databases for marine fish eDNA metabarcoding studies. This, together with additional barcoding efforts to populate reference databases, is a major milestone for making fish eDNA biomonitoring reliable and trustful.

## Data achievement statement

Raw sequencing reads and associated metadata are available on the NCBI SRA (BioProject PRJNA894161) Developed scripts and corresponding output files are available at GitHub (https://github.com/rodriguez-ezpeleta/NEA_fish_DB).

