## Supplementary material for "An automated workflow to assess completeness and curate GenBank for eDNA metabarcoding: the marine fish assemblage as case study"

AZTI, Marine Research, Basque Research and Technology Alliance (BRTA), Sukarrieta, Bizkaia, 48395, Spain

*Cristina Claver;

*Naiara Rodriguez-Ezpeleta;

### SUPPLEMENTARY MATERIAL

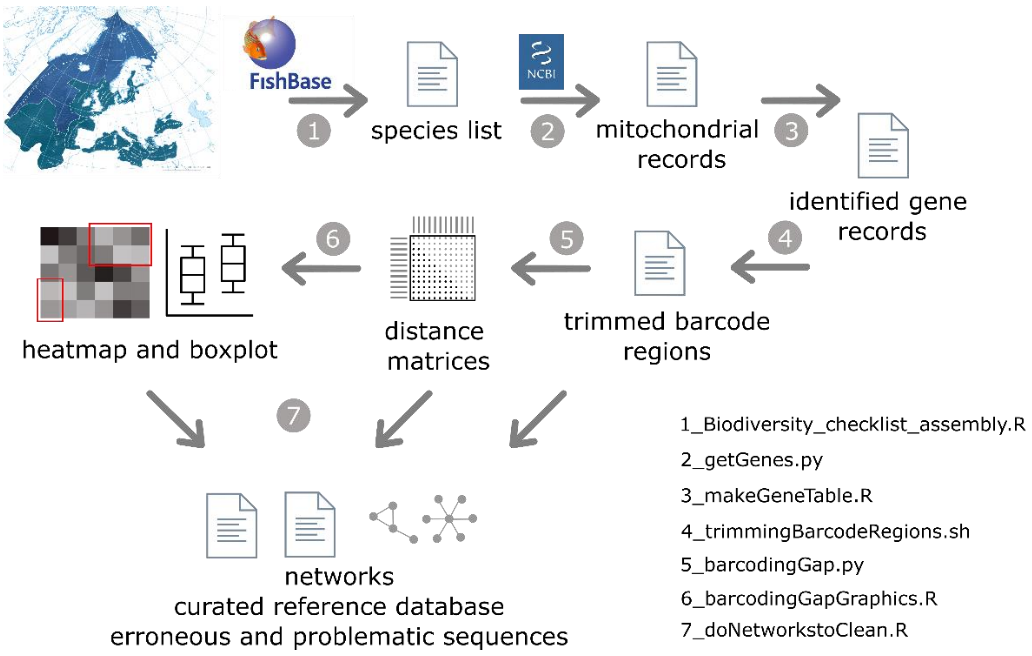


**FIGURE S1.** Schematic view of the workflow developed in this study. Numbers correspond to specific scripts.

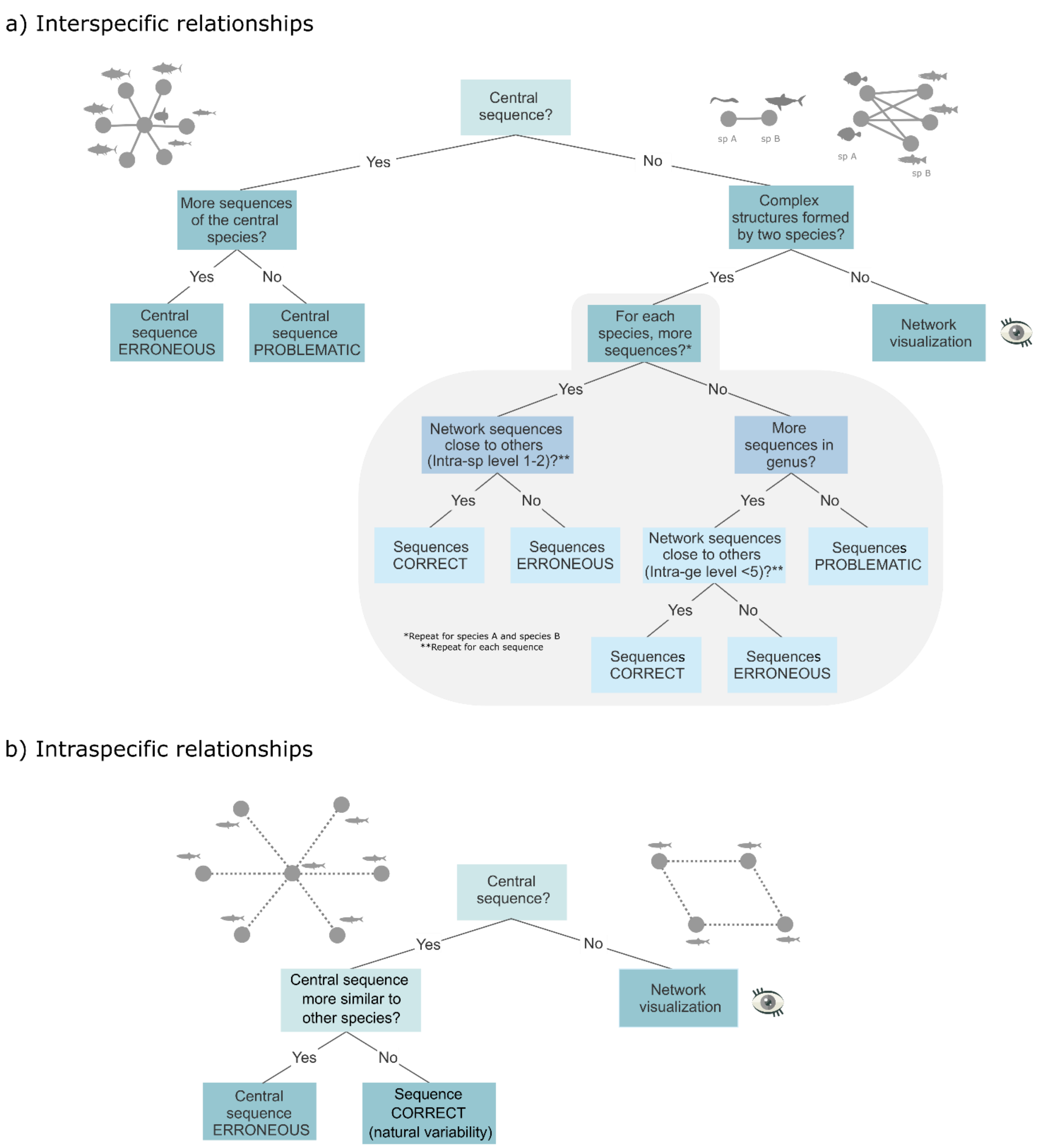


**FIGURE S2.** Decision trees of the defined criteria for spurious sequence classification based on **a)** interspecific and **b)** intraspecific relationships by network analysis.

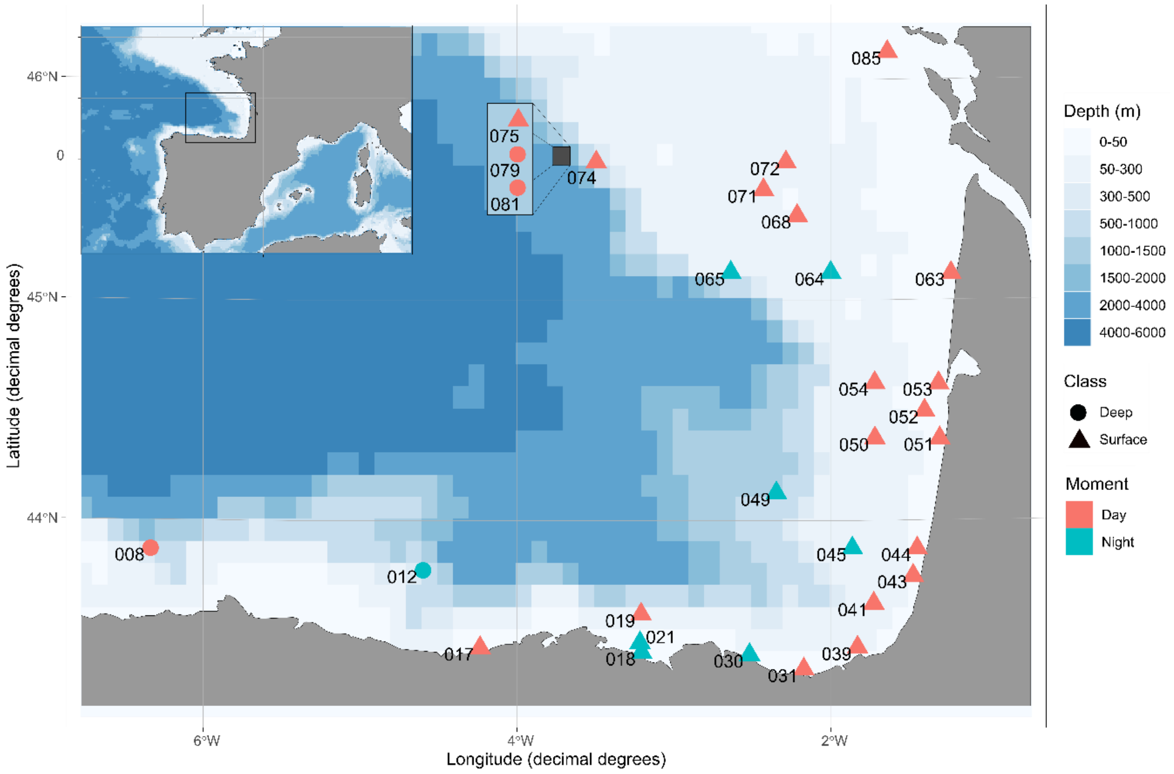


**FIGURE S3.** Map of the Bay of Biscay displaying the locations of the samples included in this study, collected in Spring 2018 on board the Ramón Margalef research vessel. Three of the samples (075, 079, 081) were collected in the same station but at different depths.

**TABLE S1.** Species from which new barcodes were generated and corresponding GenBank accession numbers.

| Species | GenBank Accession |
|---|---|
| *Argyrosomus regius* | ON000317 |
| *Benthosema glaciale* | ON000318 |
| *Ceratoscopelus maderensis* | ON000319, ON000320 |
| *Ceratoscopelus warmingii* | ON000321-ON000323 |
| *Chelidonichthys cuculus* | ON000324, ON000325 |
| *Chelidonichthys lucernus* | ON000326 |
| *Diaphus holti* | ON000327, ON000328 |
| *Labrus mixtus* | ON000329, ON000330 |
| *Lampanyctus alatus* | ON000331, ON000332 |
| *Lampanyctus pusillus* | ON000333-ON000336 |
| *Lepidorhombus whiffiagonis* | ON000337-ON000339 |
| *Lobianchia dofleini* | ON000340, ON000341 |
| *Microchirus variegatus* | ON000342 |
| *Myctophum punctatum* | ON000343-ON000345 |
| *Notolychnus valdiviae* | ON000346-ON000348 |
| *Photostomias guernei* | ON000349-ON000351 |
| *Sarda sarda* | ON000352 |
| *Umbrina canariensis* | ON000353, ON000354 |
| *Umbrina cirrosa* | ON000355 |
| *Vinciguerria attenuata* | ON000356, ON000357 |
| *Vinciguerria nimbaria* | ON000358-ON000360 |

**TABLE S2.** List of species in the European Marine regions and the number of sequences available in GenBank for each of the four genes included in this study.

*[provided as a separate file: Table_S2.txt]*

**TABLE S3**. Divergence ranges calculated for the intraspecific data for each barcode.

| Barcode | Level 1 | Level 2 | Level 3 | Level 4 | Level 5 |
|---|---|---|---|---|---|
| *folCOI* | 100-98 | 98-91 | 91-85 | 85-78 | 78-71 |
| *mlCOI* | 100-98 | 98-92 | 92-85 | 85-78 | 78-72 |
| *MiFish* | 100 | 100-93 | 93-87 | 87-80 | 80-74 |
| *teleo* | 100 | 100-96 | 96-91 | 91-87 | 87-83 |

**TABLE S4**. Erroneous and problematic sequences identified with the network analysis in *teleo* reference database.

| Species | Acc. Number | Classification |
|---|---|---|
| *Abudefduf vaigiensis* | MH248164.1 | Erroneous |
| *Alburnus alburnus* | MT410946.1 | Erroneous |
| *Anisarchus medius* | MH998262.1 | Problematic |
| *Avocettina infans* | DQ645660.1 | Erroneous |
| *Carcharodon carcharias* | AF447989.1 | Erroneous |
| *Centrolabrus exoletus* | NC_052764.1 | Problematic |
| *Cottus poecilopus* | MW448544.1 | Erroneous |
| *Holtbyrnia anomala* | KX929899.1 | Erroneous |
| *Istiophorus albicans* | AP006035.1 | Erroneous |
| *Istiophorus albicans* | NC_022478.1 | Erroneous |
| *Labrus bergylta* | MT410912.1 | Erroneous |
| *Lagocephalus laevigatus* | AY141360.1 | Erroneous |
| *Limanda limanda* | MN122886.1 | Erroneous |
| *Lutjanus argentimaculatus* | MH085613.1 | Erroneous |
| *Masturus lanceolatus* | AP006239.1 | Problematic |
| *Neogobius melanostomus* | NC_052757.1 | Problematic |
| *Osmerus eperlanus* | NC_052758.1 | Problematic |
| *Pagrus pagrus* | AY439119.1 | Erroneous |
| *Ponticola syrman* | KF415444.1 | Erroneous |
| *Pseudocarcharias kamoharai* | KM597489.1 | Erroneous |
| *Stephanolepis hispidus* | MH377775.1 | Erroneous |
| *Syngnathus acus* | JX228156.1 | Erroneous |

**TABLE S5**. Erroneous and problematic sequences identified with the network analysis in *MiFish* reference database.

| Species | Acc. Number | Classification |
|---|---|---|
| *Alburnus alburnus* | MT410946.1 | Erroneous |
| *Aluterus scriptus* | KT600933.1 | Erroneous |
| *Ammodytes tobianus* | MW818301.1 | Erroneous |
| *Bathylagichthys greyae* | LC091600.1 | Problematic |
| *Caranx crysos* | MW435597.1 | Erroneous |
| *Carcharodon carcharias* | AF447989.1 | Erroneous |
| *Cobitis taenia* | MW652799.1 | Problematic |
| *Coris julis* | MT497795.1 | Erroneous |
| *Echiodon drummondii* | KU681344.1 | Erroneous |
| *Exocoetus volitans* | LC104437.1 | Problematic |
| *Ginglymostoma cirratum* | NC_030189.1 | Problematic |
| *Hoplostethus atlanticus* | LC579079.1 | Erroneous |
| *Isurus oxyrinchus* | AF447999.1 | Erroneous |
| *Lepidorhombus whiffiagonis* | MW818389.1 | Problematic |
| *Melanolagus bericoides* | LC069498.1 | Erroneous |
| *Mugil curema* | EU715420.1 | Erroneous |
| *Nansenia groenlandica* | LC458386.1 | Problematic |
| *Neogobius melanostomus* | NC_052757.1 | Erroneous |
| *Oxynotus paradoxus* | GU130601.1 | Erroneous |
| *Ponticola kessleri* | NC_025638.1 | Problematic |
| *Pseudocaranx dentex* | MT497785.1 | Erroneous |
| *Rachycentron canadum* | FJ374798.1 | Erroneous |
| *Rhincodon typus* | KF679782.1 | Problematic |
| *Rondeletia loricata* | LC145959.1 | Erroneous |
| *Sardinella maderensis* | NC_009587.1 | Problematic |
| *Schedophilus medusophagus* | MT410878.1 | Erroneous |
| *Sphoeroides spengleri* | AY700284.1 | Erroneous |
| *Sphyrna lewini* | AF448021.1 | Erroneous |
| *Stephanolepis hispidus* | KT600974.1 | Erroneous |

**TABLE S6.** For *teleo* amplicons, number of raw sequencing reads obtained, percentage of raw reads retained after processing, and percentage of retained reads taxonomically assigned to species level using the raw and cleaned databases.

| Sample | Raw **(total)** | Retained (%) | Tax assignment Raw (%) | Tax assignment Cleaned (%) |
|---|---|---|---|---|
| 18BIO008 | 44,383 | 27 | 19 | 19 |
| 18BIO012 | 62,196 | 18 | 44 | 44 |
| 18BIO017 | 48,502 | 65 | 73 | 73 |
| 18BIO018 | 48,398 | 98 | 99 | 99 |
| 18BIO019 | 21,168 | 60 | 88 | 88 |
| 18BIO021 | 46,816 | 74 | 78 | 83 |
| 18BIO030 | 49,034 | 84 | 92 | 93 |
| 18BIO031 | 34,088 | 51 | 95 | 96 |
| 18BIO039 | 41,772 | 60 | 84 | 84 |
| 18BIO041 | 80,813 | 46 | 86 | 86 |
| 18BIO043 | 43,055 | 52 | 72 | 72 |
| 18BIO044 | 48,963 | 61 | 71 | 71 |
| 18BIO045 | 27,685 | 61 | 79 | 80 |
| 18BIO049 | 67,361 | 87 | 95 | 95 |
| 18BIO050 | 55,798 | 66 | 72 | 72 |
| 18BIO051 | 67,932 | 71 | 85 | 86 |
| 18BIO052 | 52,196 | 74 | 87 | 88 |
| 18BIO053 | 57,578 | 85 | 80 | 81 |
| 18BIO054 | 33,764 | 49 | 90 | 90 |
| 18BIO063 | 67,429 | 70 | 88 | 90 |
| 18BIO064 | 111,252 | 94 | 99 | 99 |
| 18BIO065 | 106,149 | 92 | 98 | 99 |
| 18BIO068 | 127,723 | 94 | 89 | 89 |
| 18BIO071 | 88,317 | 98 | 99 | 99 |
| 18BIO072 | 92,076 | 99 | 99 | 100 |
| 18BIO074 | 87,895 | 98 | 97 | 97 |
| 18BIO075 | 130,108 | 98 | 98 | 98 |
| 18BIO079 | 59,899 | 44 | 65 | 65 |
| 18BIO081 | 30,218 | 57 | 67 | 68 |
| 18BIO085 | 93,349 | 76 | 97 | 97 |

**TABLE S7.** For *MiFish* amplicons, number of raw sequencing reads obtained, percentage of raw reads retained after processing, and percentage of retained taxonomically assigned to species level using the raw and cleaned databases.

| Sample | Raw (%) | Retained (%) | Tax assignment Raw (%) | Tax assignment Cleaned (%) |
|---|---|---|---|---|
| 18BIO008 | 192,082 | 39 | 83 | 83 |
| 18BIO012 | 105,793 | 16 | 49 | 49 |
| 18BIO017 | 99,220 | 10 | 97 | 97 |
| 18BIO018 | 177,457 | 95 | 99 | 99 |
| 18BIO019 | 97,188 | 12 | 99 | 99 |
| 18BIO021 | 182,296 | 47 | 96 | 96 |
| 18BIO030 | 275,811 | 83 | 99 | 99 |
| 18BIO031 | 211,929 | 57 | 100 | 100 |
| 18BIO039 | 190,990 | 16 | 98 | 98 |
| 18BIO041 | 184,934 | 12 | 94 | 94 |
| 18BIO043 | 142,618 | 17 | 94 | 94 |
| 18BIO044 | 120,505 | 91 | 99 | 99 |
| 18BIO045 | 87,321 | 51 | 99 | 99 |
| 18BIO049 | 88,023 | 37 | 99 | 99 |
| 18BIO050 | 75,413 | 33 | 97 | 97 |
| 18BIO051 | 90,455 | 59 | 99 | 99 |
| 18BIO052 | 169,739 | 65 | 99 | 99 |
| 18BIO053 | 149,088 | 50 | 99 | 99 |
| 18BIO054 | 210,221 | 22 | 98 | 98 |
| 18BIO063 | 156,167 | 13 | 99 | 99 |
| 18BIO064 | 231,449 | 49 | 100 | 100 |
| 18BIO065 | 271,721 | 62 | 98 | 98 |
| 18BIO068 | 256,709 | 4 | 46 | 46 |
| 18BIO071 | 320,166 | 86 | 100 | 100 |
| 18BIO072 | 269,230 | 90 | 99 | 99 |
| 18BIO074 | 151,911 | 9 | 98 | 98 |
| 18BIO075 | 127,863 | 15 | 100 | 100 |
| 18BIO079 | 161,478 | 7 | 50 | 50 |
| 18BIO081 | 113,899 | 87 | 0 | 0 |
| 18BIO085 | 120,994 | 9 | 97 | 97 |
